# Automated detection of the HER2 gene amplification status in Fluorescence *in situ* hybridization images for the diagnostics of cancer tissues

**DOI:** 10.1101/490052

**Authors:** Falk Zakrzewski, Walter de Back, Martin Weigert, Torsten Wenke, Silke Zeugner, Robert Mantey, Christian Sperling, Katrin Friedrich, Ingo Roeder, Daniela E. Aust, Gustavo Baretton, Pia Hönscheid

## Abstract

The human epidermal growth factor receptor 2 (HER2) gene amplification status is a crucial marker for evaluating clinical therapies of breast or gastric cancer. We propose a deep learning-based pipeline for the detection, localization and classification of interphase nuclei depending on their HER2 gene amplification state in Fluorescence in situ hybridization (FISH) images. Our pipeline combines two RetinaNet-based object localization networks which are trained (1) to detect and classify interphase nuclei into distinct classes normal, low-grade and high-grade and (2) to detect and classify FISH signals into distinct classes HER2 or centromere of chromosome 17 (CEN17). By independently classifying each nucleus twice, the two-step pipeline provides both robustness and interpretability for the automated detection of the HER2 amplification status. The accuracy of our deep learning-based pipeline is on par with that of three pathologists and FISH images on a set of 57 validation images containing several hundreds of nuclei are accurately classified. The automatic pipeline is a first step towards assisting pathologists in evaluating the HER2 status of tumors using FISH images, for analyzing FISH images in retrospective studies, and for optimizing the documentation of each tumor sample by automatically annotating and reporting of the HER2 gene amplification specificities.

## Background

The human epidermal growth factor receptor 2 (HER2) gene, also designated ERBB2 gene for the v-erb-b2 erythroblastic leukemia viral oncogene homolog 2, encodes a member of the epidermal growth factor receptor family of receptor tyrosine kinases. Amplification of the HER2 gene is the primary mechanism of HER2 overexpression in tumors^1^. HER2 amplification occurs before HER2 protein overexpression and monitoring of the tumor HER2 gene amplification status has therefore become routine in breast cancer^2–4^ surveillance. In around 25% of breast cancers a positive HER2 status is associated with poorer prognosis, more aggressive disease, and an increased risk of disease recurrence^2,5–7^. Application of HER2-directed therapies such as treatment with anti-HER2 antibodies, e.g. trastuzumab, depends on the detection of HER2 gene amplification and increases overall survival of individuals suffering from HER2 positive breast cancer^2,6–10^. In addition to breast cancer, HER2 status testing is also applied in gastric cancers as trastuzumab is similarly effective in prolonging survival in HER2 positive carcinoma of the gastric and of the gastroesophageal junction^2,11^.

HER2 testing is commonly carried out by immunohistochemistry (IHC), chromogenic *in situ* hybridization (CISH), silver-enhanced *in situ* hybridization (SISH) or Fluorescence *in situ* hybridization (FISH). In interphase nuclei of investigated tumor material, HER2 gene amplification testing is preferentially conducted via FISH^12^. In FISH analysis a HER2 positive state is defined when a HER2/CEN17 ratio of more than 2.2 is detectable, where CEN17 is a centromeric probe for the centromere of chromosome 17 on which the HER2 gene resides. Negative HER2 FISH amplification is defined as HER2/CEN17 ratio of less than 1.8^12^. If an internal control probe such as CEN17 is not available, HER2 positive FISH is defined when above six HER2 genes are detectable per interphase nucleus while the equivocal range is defined with an average copy number of four to six HER2 genes per nucleus. Normal nuclei harbor two or fewer HER2 genes^13^.

In clinical practice, the analysis is carried out by a pathologist via observation of the FISH slide using a fluorescence microscope. The decision making relies on the individual expert knowledge of the pathologist and is dependent on standardization of the methodology, lab-dependent routines, and finally on the quality of the FISH images, which is influenced by e.g. background signal, artifacts, tissue quality, and other acquisition-dependent parameters. Pathologists analyze the HER2 gene amplification status of a tumor sample via evaluation in comparison to control samples. Testing criteria define HER2 positive status when (on observing within an area of tumor that amounts to > 10% of contiguous and homogeneous tumor nuclei) there is evidence of HER2 gene amplification based on counting at least 20 nuclei within this area^14^. By counting and classification of at least 20 interphase nuclei from different areas of the FISH slides a diagnostic decision is possible regarding a positive or negative state of HER2 gene amplification and its HER2 grade (low or high). The diagnostic relies on ratios of HER2 to CEN17 signals per nuclei on which the subsequent classification of the corresponding tumor sample is conducted.

While many classical methods exist for automatic extraction of features from microscopic images such as spot detection and counting^15,16^, during the last years an increasing number of deep learning based applications for classification tasks of pathological microscopic images were developed and successfully conducted on a wide field of applications^17^. There, image classification tasks commonly involve the application of Convolutional Neural Networks (CNNs) that rely on a stack of convolutional and non-linear transformations of the input data to create high-level abstraction classification^18^. Deep learning approaches such as CNNs have been already performed on pathology image classification, tumor classification, on imaging mass spectrometry data^16^, in the identification of metastatic cancer areas^17^, and annotation of pathological images^19^. In the context of FISH images, CNNs have been used for segmentation of chromosomes in multicolor FISH images^20^ and for detecting and counting of fluorescence signal in nuclei (SpotLearn)^21^. SpotLearn includes two supervised machine learning-based analysis workflows for high-accuracy detection of FISH signals from images with three separate fluorescence microscopy channels^21^. However, in the certified routine diagnostic workflow established in our Institute of Pathology, FISH signals are captured using a graded filter and the different HER2 gene and CEN17 signals are recorded in one step. Consequently, the resulting single-channel images cannot be distinguished by SpotLearn. Moreover, whereas SpotLearn detects spots of fluorescence, called FISH signals in our study, via segmentation, we aim to both localize and classify nuclei and FISH signals without the need for segmentation and additionally provide a detailed report on the HER2 amplification status in the sample.

To address these issues, we developed a pipeline for automatic detection of HER2 amplification status in FISH images based on RetinaNet, a state-of-the-art CNN for object localization. The pipeline consists of two independently trained and validated object localization networks. In the first step, nuclei are localized and classified as low, normal or high grade in a whole FISH image. Subsequently, for each detected nucleus, a second network localizes and classifies each individual fluorescence signal as HER2 or CEN17 from which HER2/CEN17 ratios are calculated.

We demonstrate that this two-step process provides a per-nucleus classification accuracy of the amplification status that is on par with the interrater agreement between three pathologists. Moreover, the classification accuracy of whole FISH images, achieved by combining the results of the two detection networks, was found to be in nearly perfect agreement with the team of pathologists. By classifying each nucleus twice our pipeline intrinsically provides double reading to expose prediction uncertainty which is essential in clinical applications. Additionally, our detection system yields interpretable results by providing a detailed report on the amplification status of every nucleus in the FISH image. This allows pathologists to understand the decision of our deep learning system, which is a prerequisite for the critical questioning of that decision and for the manual reassessment of questionable or uncertain cases. In clinical practice, our deep learning system could provide an assisting platform for pathologists in the daily routine diagnosis of HER2 amplification status detection in FISH image analysis of breast and gastric cancer. In addition, all nuclei of a FISH can be analyzed at once, enabling HER2 amplification state recognition based on whole FISH slide annotation.

## Material and Methods

### Preparation of slides, Fluorescence in situ hybridization (FISH) and image capturing

Formalin-fixed Paraffin-embedded (FFPE) cancer tissue was delivered from clinical institutions from all over Germany. FFPE tissue was cut into small pieces (2μm) on a slide and dehydrated using first a xylene washing step subsequent flowed by a series of ethanol steps (100%, 96%, 70%). After drying the slide at room temperature slides were incubated with sodium thiocyanate followed by a wash step using distilled water. Subsequently, slides were incubated with pepsin and hydrochloric acid, washed using distilled water and dried at room temperature. Probes (PathVysion HER-2 DNA Probe Kit II, Abbott Inc.) were hybridized at 37°C in a wet chamber overnight. Washing of slides was performed in 2x saline-sodium citrat**e**(SSC) buffer and DAPI counterstaining was conducted. Images were taken using fluorescence microscope (Axioskop 2, Zeiss Inc.) using a graded filter (Filter Set 23 (488023-0000-000), emission: 515-530 nm + 580-630 nm, Zeiss Inc.), recording HER2 gene signals, CEN17 signals and a small subset of DAPI signals at once. Images were captured at a magnification of 40x and processed using the Image-Pro 6.0 software (Media Cybernetics) and saved in JPEG file format with a size of 1200 × 1600 pixel.

### Convolutional neural network architecture

Our pipeline consists of two convolutional neural networks (CNN) for object localization. The “nucleus detector network” takes the whole FISH image as input and localizes nuclei while simultaneously classifying them into *high*, *normal*, or *low grade* (**Figure 1A**). The “signal detector network” takes cropped image-regions around the already detected nuclei as input and localizes the individual spot-like fluorescence signals therein and classifies them into HER2 or CEN17 gene signals **(Figure 1B)**. Both detector networks have an identical architecture based on RetinaNet (**Figure 1C)** and have identical training procedures.

**Figure 1.**
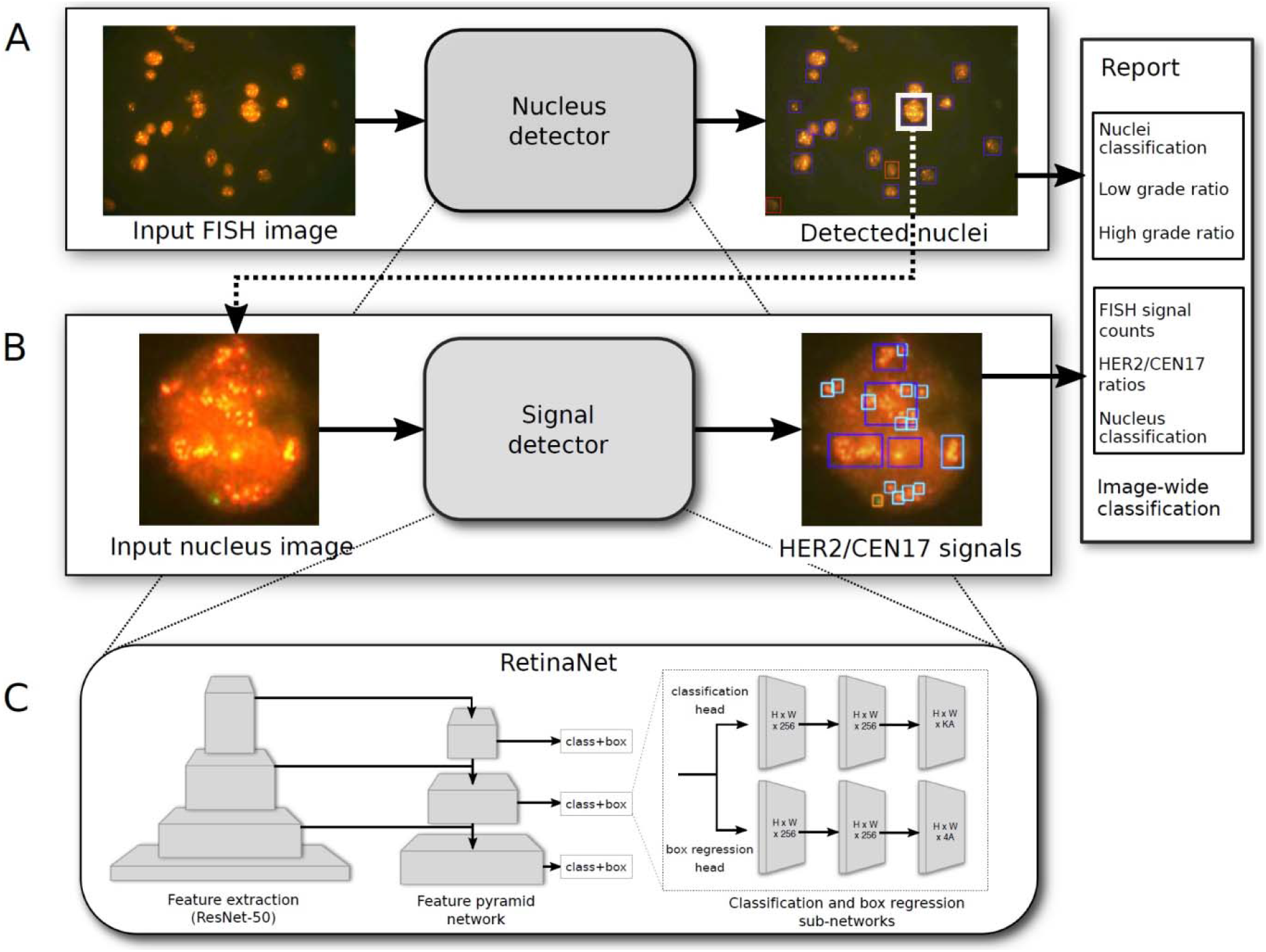
Illustration of the two-stage deep learning detection system of the HER2 gene amplification stage in FISH images from breast cancer samples. **(A)** The nucleus detector network takes whole FISH images as input and outputs the localization and classification for all detected nuclei. **(B)** The signal detector network subsequently takes each detected nucleus and localizes and classifies individual FISH signals. The output of both networks is post-processed by calculation of the low/high grade ratios and HER2/CEN17 ratios, and an image-wide classification prediction is computed and reported. **(C)** Both detectors are based on RetinaNet which consists of a ResNet50 feature extraction network, a feature pyramid network and two fully convolutional classification and box regression networks for every level of the feature pyramid.

RetinaNet^22^ is a state-of-the-art convolutional neural network (CNN) for object localization. Its architecture consists of three key parts: First, a standard feedforward CNN with convolutional and pooling layers (ResNet50^23^) is used as backbone network to perform general feature extraction (**Figure 1C**, **left**). Second, the resulting feature maps are used in a Feature Pyramid Network (FPN)^24^ that provides semantically rich feature maps at various spatial resolutions (**Figure 1C**, **center**) by combining the up sampled features of low resolution but high semantic content with the high resolution feature maps from the corresponding spatial scale. Third, two fully convolutional heads are attached at each level of the FPN that compute, for each anchor box at every spatial location, the offsets of the bounding boxes for detected objects (regression head) and the class probabilities for each object (classification head) (**Figure 1C**, **right**). The large number of boxes predicted by RetinaNet improves the accuracy compared to one-step detector networks such as YOLO^25,26^ and SSD^27^, particularly for small objects. However, it also causes an extreme class imbalance as most predicted boxes are easy-to-classify background samples. In order to avoid the loss being dominated by these background samples, RetinaNet replaces the typically used cross-entropy loss with a dedicated loss function (focal loss) that increases the contribution from hard examples while decreasing the contribution of uninformative background samples.

In this work, we used the implementation of RetinaNet provided by Fizyr^28^ with a ResNet50 backbone provided by the Broad Institute. All networks are implemented within the Keras^29^ framework with Tensorflow^30^ as backend.

### Image annotation

FISH images of high quality were selected from documented images of breast cancer FISH diagnostics on the HER2 gene amplification status from the years 2015-2018 harbored at the Institute of Pathology at the clinical campus of Carl Gustav Carus Hospital of TU Dresden. FISH images (n=299) were manually annotated by providing bounding boxes and class labels for every nucleus in these images. Nuclei were categories into five classes: *low*, *normal*, *high grade*, *uncertain,* and *artefact*. Additionally, images of individual nuclei (n=300) were manually annotated with bounding boxes and class labels for each individual FISH signal, classified as CEN17, HER2 or HER2 cluster. The latter class was introduced to represent a cluster of HER2 signals where individual signals could not be distinguished. Annotation was performed manually by a pathologist using LabelImg^31^. Training and validation (randomly chosen 10% of all images) was performed on each step, respectively. An overview of the training data set is given in **Table 1** for the nucleus detector and in **Table 2** for the signal detector.

**Table 1.** Details on training FISH images.

|  | # nuclei |  |  |  |  |
| --- | --- | --- | --- | --- | --- |
| # images | normal | low-grade | high-grade | # uncertain | # artifact |
| 299 | 626 | 782 | 1,760 | 2,037 | 1,050 |

**Table 2.** Details on training interphase nuclei.

|  | # FISH signals |  |  |
| --- | --- | --- | --- |
| # images | CEN17 | HER2 | HER2 cluster |
| 301 | 512 | 1,552 | 441 |

### Training procedure

Except for the input data and annotation, identical training procedures, loss functions and hyperparameters were used for training of both networks. Image data (including ground-truth bounding boxes) was augmented using rotations, random crops, translations, shearing, scaling, and horizontal and vertical flips using augmentation implementation provided by the Keras RetinaNet package. We used focal loss^24^ for the classification, and a smooth L1 loss for bounding box regression. For optimization we used adaptive moment estimation (Adam)^28^ with a fixed learning rate of 10^−4^ and a batch size of 1 due to GPU memory limitations. Both networks were trained for 50 epochs without pre-training on a single NVIDIA GPU (GeForce 1080Ti).

### Post processing

To convert the localization and classification results from the nucleus detector network and the signal detector network into nucleus-level and image-level predictions, we implemented detector-specific post processing steps as part of the pipeline. Concretely, the results of the nucleus detector are used to calculate two ratios: ratio-1 is the number of low-grade nuclei divided by the number of all detected nuclei; ratio-2 is the number of high-grade nuclei divided by the number of all detected nuclei. A FISH image is defined to be low-grade when ratio-1 is at least 0.2 while a FISH image is classified to be high-grade when ratio-2 is at least 0.4. These thresholds can be modified by the pathologist according to individual specificities and criteria.

Similarly, analogous ratios are computed from the results of the signal detector network. Specifically, the HER2/CEN17 ratio is calculated as the ratio of the number of CEN17 signals to the number of HER2 signals per nucleus. In case no HER2 signals were detected, the ratio is set to 1. In case of detection of a HER2 cluster, the ratio is set to a high value (HER2/CEN17=10). In case no CEN17 signal was detected, the nucleus is classified as artefact. To classify the FISH image based on the HER2/CEN17 ratios for each nucleus, the average of the HER2/CEN17 ratios is calculated. A HER2/CEN17 ratio greater than 1.5 and lower than 6.0 indicates a low-grade status of the FISH image. A value greater than 6.0 indicates a high-grade status of the FISH image. These thresholds can be modified by the pathologist according to individual specificities and criteria. The localization and classification results, as well as the calculated ratios for both detector networks were stored in a text file alongside the annotated images. From these files, a detailed report was generated for each FISH image describing the final image-wide classification as well as the nucleus- and signal-level classification details on which this decision is based.

## Results

To enable an automated detection service for FISH image samples, we developed a pipeline based on convolutional neural networks (CNN) for object detection, and trained it on annotated FFPE breast cancer FISH image samples originating from routine diagnostics. The data set used in this retrospective study was obtained using a certified FISH protocol that is used routinely on the daily diagnostic practice for breast and gastric cancer patients which has been in use since 16 years. Our deep learning pipeline consists of two RetinaNet^22^-based object localization networks that successively process a single FISH image. First, the nucleus-detector network localizes and classifies the amplification status of each nucleus in the FISH image into low, normal or high grade (**Fig. 1A**). Subsequently, the signal-detector network processes cropped images of each nucleus detected in the first step, and localizes and classifies every HER2 and CEN17 signal in these image, from which a HER2/CEN17 ratio is calculated (**Fig. 1B**). In this fashion, every nucleus in the FISH image is classified twice by two independently trained CNN models. From these nucleus-level classifications, proportions between low and high grade nuclei are calculated that result in a final image-wide classification into low, normal or high grade amplification status. This decision-making process was designed to be analogous to the assessment made by human pathologists where in a first step nuclei are image-wide localized and classified and secondly a confirmation of the classification is applied on the basis of HER2/CEN17 ratios for each nucleus (**Fig. 1**).

### Nucleus detector: Detection and classification of nuclei in FISH images

Training of the nucleus-detector network was performed on manually labelled FISH images (n=299) containing several thousands (n~7,000) of high-grade, low-grade and normal nuclei, as well as uncertain cases and artifacts (as described in materials and methods) (**Tab. 1**).

To validate the applicability and reliability of the nucleus-detector network in routine diagnostics, 57 high quality FISH images were subject to image-wide nuclei detection and classification and compared to the annotation by three pathologists. A total of 1,183 nuclei were independently evaluated by a team of three pathologists on the one hand and the nucleus detector in the other hand. Both pathologists and the deep object detector network were tasked to localize and classify each nucleus in the 57 FISH images as unidentifiable (incl. artifacts and uncertain cases), low-grade, normal, or high-grade. The classification results were collected as confusion matrices from which we calculated the interrater agreement between the nucleus detector and the pathologists in terms of a weighted Cohen’s kappa coefficient κ^32^ reflecting the agreement between independent observers and the ordinal nature of the classes. In absence of an unambiguous ground truth data set, we calculate the arithmetic mean of the agreement coefficients κ to reflect the performance of the nucleus detector compared to three pathologists.

As shown in **Figure 2A**, we find that the nucleus detector obtains a mean κ_ND_=0.648, representing a substantial level of agreement^32^ between detector and pathologists. To compare this result to the agreement among the pathologists themselves, we calculated the agreement between the annotations obtained by all three pathologist using the average pairwise Cohen’s kappa’s (also known as Light’s kappa)^32^, resulting in κ_patho_=0.643. This demonstrates that there is a similar classification reliability among human pathologists, reflecting the inherent ambiguities in reading FISH images. As indicated in the accuracies displayed in Figure 2B and as broken down in the confusion matrices in **Figure 2C**, these ambiguities mainly affect the classification of low and normal-grade nuclei, while high-grade and undefined nuclei are classified more reliably. Importantly, since these difficulties affect both human readers and our automated pipeline and, the agreement between nucleus detector and pathologists is on par with the agreement within the team of pathologists.

**Figure 2.**
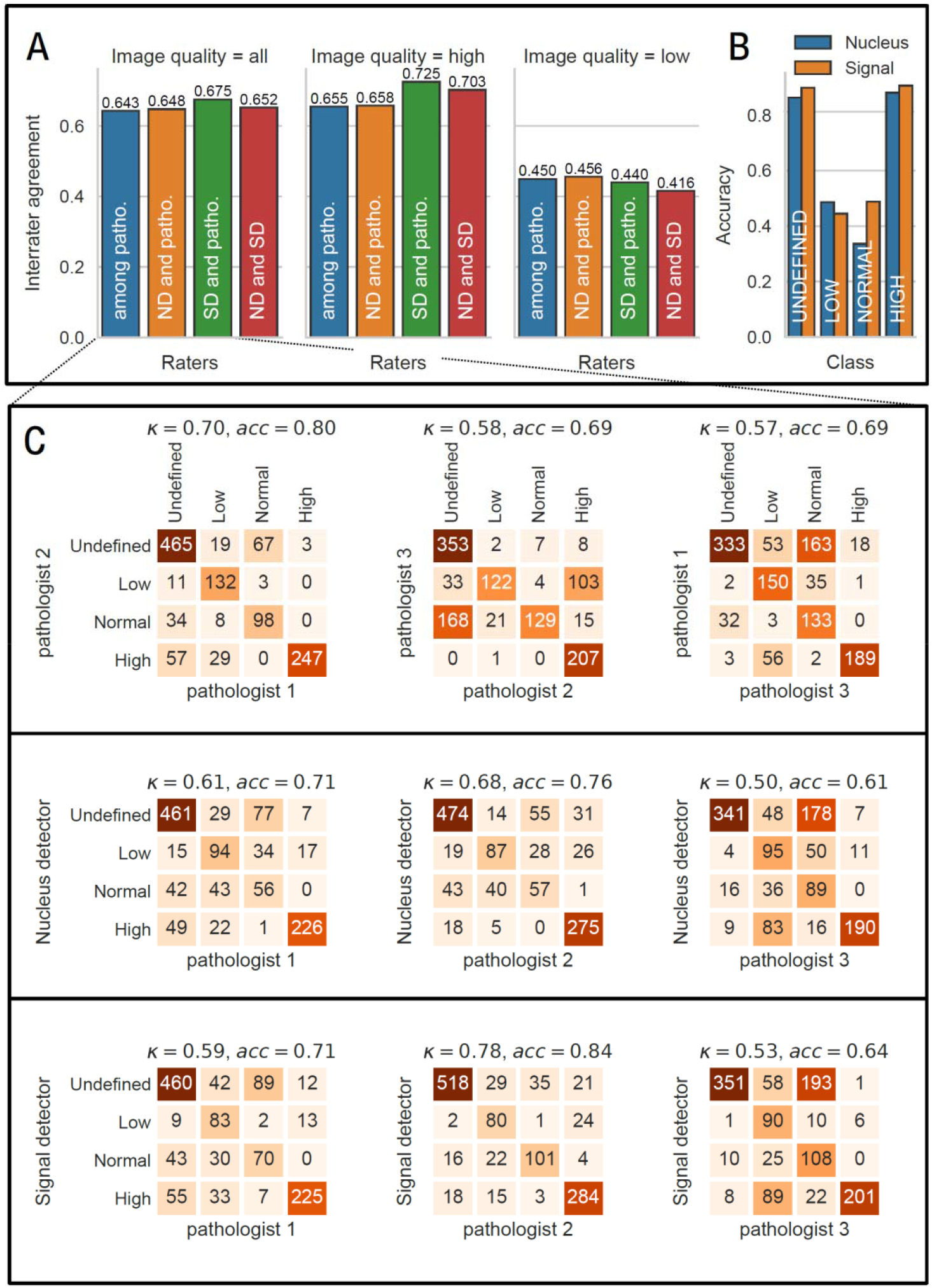
The predication of the two-stage deep learning-based detection system for the HER2 gene amplification stage is in par with pathologists. **(A)** Interrater agreement (Light’s Kappa κ) among a team of three pathologists and between the team of pathologists and the nucleus detector (ND) and signals detector (SD), respectively, for all images, for high quality images and for low quality images. **(B)** Nucleus class-specific accuracies between the team of pathologists and the nucleus and diagonal detector. **(C)** Confusion matrices for all images with respect to the classification of nuclei among the group of the three pathologists, between the three pathologists and the nucleus detector and between the three pathologists and the signal detector, respectively. Light’s Kappa κ and accuracies are shown above each matrix.

The nuclei in FISH images from our routine diagnostics in this retrospective study can be of reduced quality compared to *up*-*to*-*date* fluorescence images as they have to be prepared under time limitation and a standardization procedure. Background noise, an increased number of artifacts and large differences in the number and shape of nuclei as well as overlapping nuclei all influence the image quality of the captured nuclei. In addition, the quality depends on the input tumor material and available tissue type for analysis. To test the robustness of the nucleus detector, we manually subdivided the nuclei from our investigated FISH images into the two groups “high quality” and “low quality” nuclei. Nuclei in the “high quality” group (n=641) are characterized by clearly differentiable HER2 and CEN17 signals and exhibit a uniform and regular nucleus shape without overlapping by further nuclei. In contrast, nuclei in the “low quality” group (n=542) show blurring of FISH signals, overlapping by further nuclei, very weak FISH signals or signal artifacts which made it difficult to adequately detect the signals. As shown in **Figure 2A**, the image quality has substantial impact on the classification performance assessed by the interrater agreement. Cohen kappa of the nucleus detector is reduced from substantial agreement (κ=0.648) for high quality nuclei to moderate agreement (κ=0.458) in the case of low quality nuclei. Correspondingly, a similar decrease (from κ=0.643 to κ=0.450) can be observed in the agreement among the team of pathologists, again demonstrating the similarity in performance of our pipeline.

### Signal Detector: Counting the HER2 and CEN17 FISH signals per nucleus and detection of the image-wide HER2/CEN17 ratio

In order to increase the classification accuracy and to validate and control the nucleus-based classification performed by the nucleus detector, we trained a second RetinaNet to localize and classify the individual HER2 and CEN17 signals in each detected nucleus. It acts as a control mechanism comparable to a second opinion and provides additional detailed source of information on the number of HER2 and CEN17 signals per nucleus.

The HER2 FISH signal detection class was split into two subclasses: HER2 single signals and HER2 clusters. A HER2 cluster represents a region of the nucleus which is characterized by a high density of adjacent HER2 signals which often cannot be well distinguished into the underlying single signals and hence appeared as an accumulation. Training was performed on several thousands (n~3,000; HER2-Cluster were counted as n=1 although they contain many single HER2 gene signals) of HER2 and CEN17 signals from the ~300 randomly selected nuclei from the ~300 training FISH images. Apart from the different input images and annotations, the training procedure of the signal detector was identical to that described above for the nucleus detector (see Materials and Methods).

The signal detector predicts a bounding box and classifies each individual HER2 single signal, HER2 cluster and CEN17 signal. The boxes are counted and the ratio of HER2/CEN17 signals is calculated per nucleus. Each nucleus is classified based on this ratio as described above (see **Post-processing** in **Materials and Methods**). The average and image-wide HER2/CEN17 ratio was calculated on the basis of all detected nuclei harboring CEN17 and HER2 signals. This quantity was used to decide on the image-wide HER2 gene amplification status of the corresponding FISH image.

To measure the performance of the signal detector at the level of the detection of individual FISH signals, 50 randomly selected nuclei were analyzed and compared to the manual annotation by a pathologist. In six cases a different classification was revealed (**Tab. 2**). In three of the six cases, a normal nucleus was classified via the signal detector while the pathologist detected a low-grade nucleus which was caused due to missed HER2 single signal detection via the signal detector. In two of the six cases the signal detector identified a high-grade nucleus while the pathologist classified these nuclei as normal. The reason was that signal detector detected a HER2 signal as HER2 cluster because of strong blurring of the single HER2 signal mimicking a HER2 cluster. In one of the six cases, a classification via the signal detector was not possible although the same number of HER2 signals and CEN17 signals was found in comparison to the pathologist. However, because only one HER2 signal was identified, the signal detector classified the nucleus as “uncertain”.

To validate the applicability and reliability of the signal detector at the level of nucleus classification, the same 57 test FISH images were subject to nuclei detection and classification and compared to the team of pathologists (n=3). The comparison was also conducted to “high quality” and “low quality” nuclei as previously done for the nucleus detector to test the robustness on nuclei images of lower quality. We find an agreement value of κ=0.675, demonstrating a substantial agreement between our deep learning system and human pathologists (**Fig. 2B**). This is a slightly higher level of agreement than obtained by the nucleus detector and also higher than the agreement among pathologists. For low quality nuclei, we find only a moderate agreement (κ=0.440), which is similar to the performance of the nucleus detector. For nuclei recorded at high quality, we find a classification agreement of κ=0.725, representing a good agreement with the pathologists. Nevertheless, visual inspection showed that a minor number of HER2 double or triple signals in very close vicinity were annotated as HER2 cluster leading to the wrong overall assessment that a high-grade nucleus occurred.

### Comparison of nucleus and signal detector networks

The accuracy of both deep learning networks was compared regarding the image-wide detection and classification of nuclei in the 57 test FISH images (**Suppl. Tab. 1**). Similar to the nucleus detector, we found for the signal detector that the detection and classification performance differs between images. Interestingly, however, several images where the nucleus detector performed poorly were well-classified by the signal detector and vice versa, indicating these two approaches are complementary and are best used in combination (**Suppl. Tab. 1**). While for most images, however, the performance of both networks is similar (**Fig. 3A**), a few images show larger differences in their accuracies. Four example images are depicted where (1) the accuracy was nearly 100% for both detector networks (**Fig. 3**, **image 35**), (2) the accuracy was lowest for the signal but higher in the nucleus detector (**Fig. 3**, **image 8**), (3) the accuracy was lower for the nucleus detector compared to the signal detector (**Fig. 3**, **image 14**) and (4) the accuracy was low for both detectors (**Fig. 3**, **image 25**). Nuclei in the FISH images are marked with a red arrow where a different classification was obtained by both detectors (**Fig. 3C**). In image **35** nuclei are clearly distinguishable and show massive amplification of the HER2 gene, which can be easily and clearly detected by both detectors. Therefore, no differences in the nuclei classification were detected. In image **25** the performance of the two detectors is equally low due to the general low quality of many nuclei occurring in the image. In addition, the overall number of classifiable nuclei in the image is low so that the influence of the “low quality” nuclei on the image-wide classification is higher. Reasons for different accuracy between the nucleus and signal detectors in images **8** and **14** may be due to weak and blurring FISH signals not seen by the signal detector and/or the interpretation of very adjacent located HER2 gene signals as HER2 cluster by the signal detector leading to false classification of the corresponding nucleus. More precisely, **Figure 4** shows selected and representative examples on three cases for a same and three cases for a different classification between both detectors. The three nuclei in the left column (**Fig. 4A-C**) were classified identically by both networks and the classification corresponds to those of the pathologist, providing stronger confidence in the correct classification. The three nuclei in the right column, however, were classified differently (**Fig. 4D-F**). In the first case (**Fig. 4D**) the signal detector detected three HER2 signals in close vicinity as a single HER2 cluster, leading to a misclassification as high-grade nucleus while the nucleus detector correctly classified this nucleus as low-grade. In the second case (**Fig. 4E**), the signal detector missed the detection of HER2 and CEN17 signals, presumably due to overexposure, and therefore a misclassification of the nucleus as normal was conducted. The nucleus detector correctly classified the nucleus as low-grade. Finally, in **Figure 4F**, the signal detector correctly detected all signals but classified the nucleus as normal, the nucleus detector classified the nucleus as uncertain, and the pathologist’s classification was low-grade.

**Figure 3.**
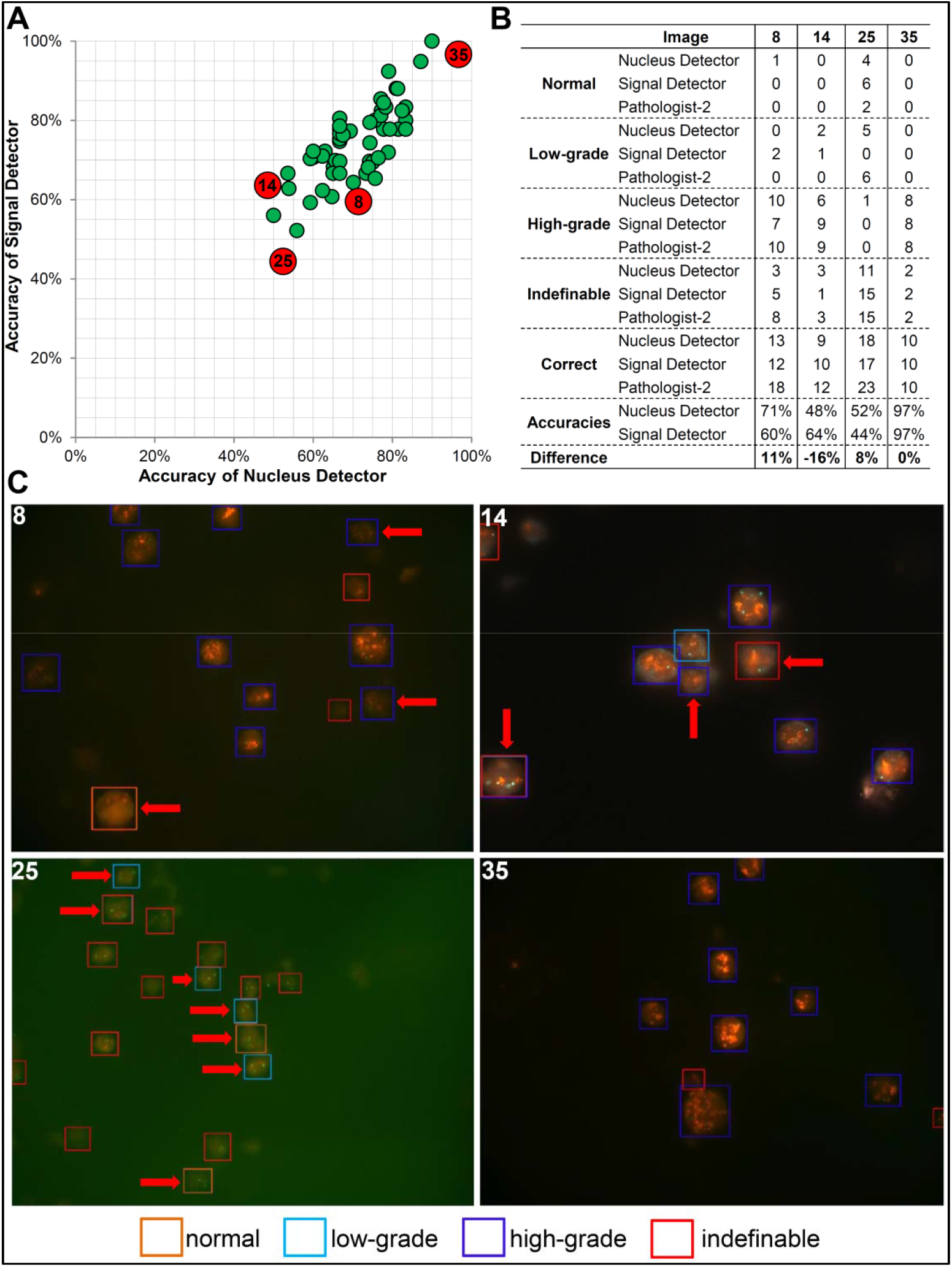
Comparison of the accuracy of nucleus detector and signal detector on the image-wide nucleus detection and classification. **(A)** The accuracy in classifying the detected nuclei was compared among the 57 validation FISH images. Exemplarily, the four most interesting images are depicted where (1) the accuracy was nearly 100% in both steps (image 35), (2) the accuracy was low in both steps (image 25), (3) nucleus detector had a better accuracy than signal detector (image 8) or (4) vice versa (image 14). **(B)** Detailed overview about the classification of the nuclei in these four FISH images and **(C)** Visualization of the classification of nucleus detector. Nuclei with a differing classification by signal detector are marked with a red arrow.

**Figure 4.**
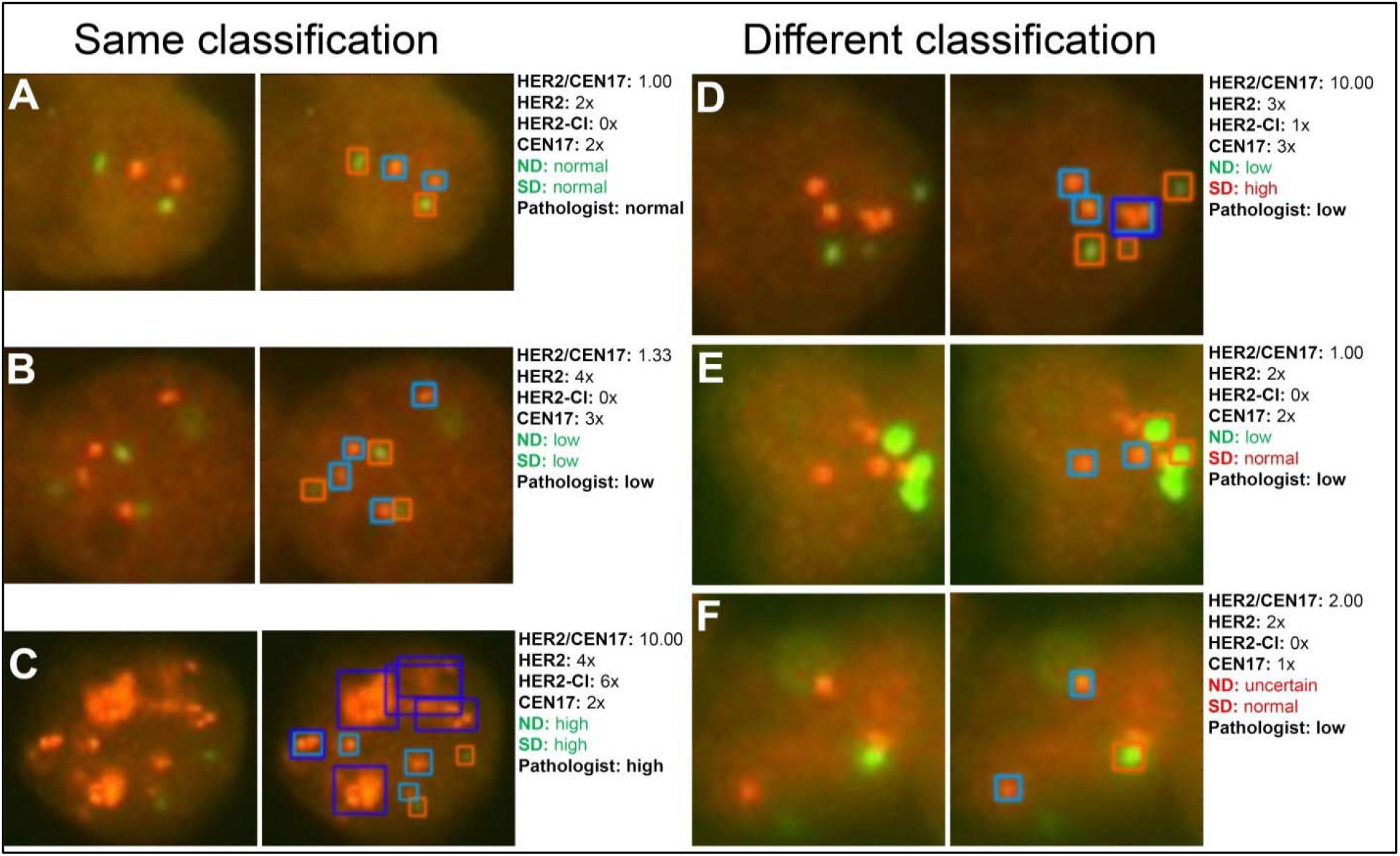
Application of the deep learning-based system on six interphase nuclei detection. The signal detector (SD) detects FISH signals in each of the separated and pre-classified nuclei from the nucleus detector (ND) and classifies the signals into HER2 (light blue framed boxes), HER2 cluster (dark blue framed boxes) and CEN17 (red framed boxes) signals. Exemplarily three nuclei are shown: **(A-C)** The classification was confirmed. **(D)** The classification was not confirmed due to a misinterpretation of three very adjacent HER2 gene single signals as HER2 cluster. **(E)** The classification was not confirmed because two HER2 gene signals were not detected by the signal detector. **(F)** The classification was not confirmed although all FISH signals were detected by signal detector because the classification was defined to be unclassifiable if only one CEN17 signals occurs.

### Automated classification of high quality FISH images into normal, low- and high-grade

Our nuclei detection and classification system relies on the combination of the two steps performed by the nucleus detectors and the signal detector enabling a final decision on the HER2 gene amplification status with HER2 and CEN17 FISH signal counting of the whole FISH image. The decision relies on ratios being calculated in both detector steps. In the nucleus detector, the ratio-1 and the ratio-2 (ranging from 0 to 1, respectively) are calculated and indicate the relative number of low grade nuclei (ratio-1) and high grade nuclei (ratio-2), respectively. A low-grade or high-grade stage is indicated by ratio-1 greater or equal to 0.2 or by ratio-2 greater than 0.4, respectively. Both thresholds are modifiable with respect to the pathologist’s specified criteria. In the signal detector, an image-wide HER2/CEN17 ratio is calculated as average value among all nuclei-specific HER2/CEN17 ratios of classifiable nuclei. A HER2/CEN17 ratio greater than 1.5 and lower than 6.0 indicates a low-grade status of the FISH image. A value greater than 6.0 indicates a high-grade status of the FISH image. The image-wide classification of the HER2 gene amplification status showed identical results by our pipeline and the team of pathologists for 55 of the 57 (96%) test FISH images, showing excellent agreement between our automated pipeline and the team of pathologists (**Tab. 3**). In two of the 57 images a different classification was observed. In one of the two cases, the nucleus detector classified a low-grade FISH image while the signal detector had a tendency towards a high-grade image. In the second case, the nucleus detector classified the image to be low-grade while the signal detector classified it as high-grade due to a misclassification of one normal nucleus as high-grade nucleus.

**Table 3.**
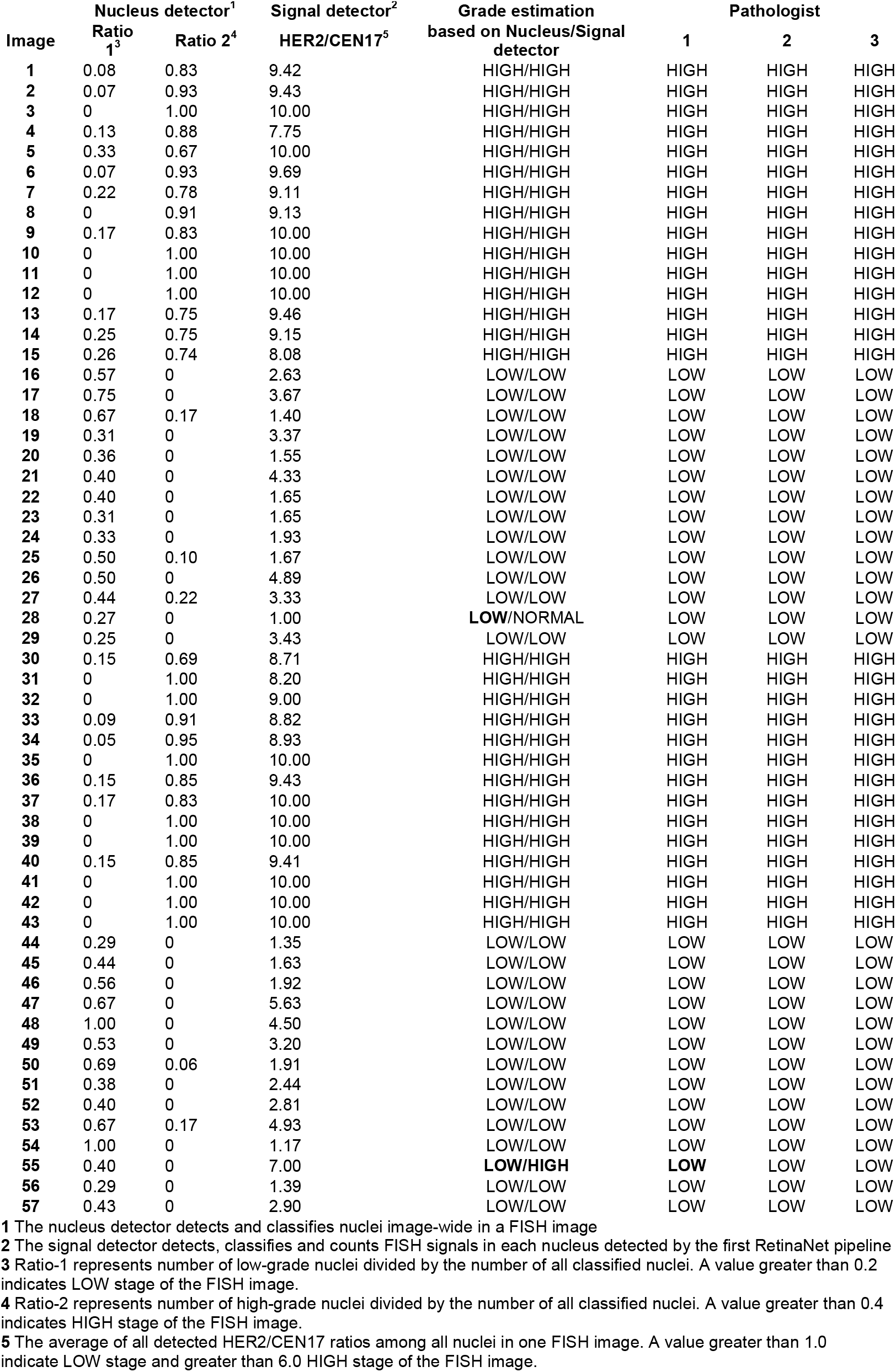
Detection of the HER2 gene amplification status in 57 validation FISH images.

| Image | Nucleus detector<sup>1</sup> |  | Signal detector<sup>2</sup> | Grade estimation | Pathologist |  |  |
| --- | --- | --- | --- | --- | --- | --- | --- |
|  | Ratio 1<sup>3</sup> | Ratio 2<sup>4</sup> | HER2/CEN17<sup>5</sup> | based on Nucleus/Signal detector | 1 | 2 | 3 |
| 1 | 0.08 | 0.83 | 9.42 | HIGH/HIGH | HIGH | HIGH | HIGH |
| 2 | 0.07 | 0.93 | 9.43 | HIGH/HIGH | HIGH | HIGH | HIGH |
| 3 | 0 | 1.00 | 10.00 | HIGH/HIGH | HIGH | HIGH | HIGH |
| 4 | 0.13 | 0.88 | 7.75 | HIGH/HIGH | HIGH | HIGH | HIGH |
| 5 | 0.33 | 0.67 | 10.00 | HIGH/HIGH | HIGH | HIGH | HIGH |
| 6 | 0.07 | 0.93 | 9.69 | HIGH/HIGH | HIGH | HIGH | HIGH |
| 7 | 0.22 | 0.78 | 9.11 | HIGH/HIGH | HIGH | HIGH | HIGH |
| 8 | 0 | 0.91 | 9.13 | HIGH/HIGH | HIGH | HIGH | HIGH |
| 9 | 0.17 | 0.83 | 10.00 | HIGH/HIGH | HIGH | HIGH | HIGH |
| 10 | 0 | 1.00 | 10.00 | HIGH/HIGH | HIGH | HIGH | HIGH |
| 11 | 0 | 1.00 | 10.00 | HIGH/HIGH | HIGH | HIGH | HIGH |
| 12 | 0 | 1.00 | 10.00 | HIGH/HIGH | HIGH | HIGH | HIGH |
| 13 | 0.17 | 0.75 | 9.46 | HIGH/HIGH | HIGH | HIGH | HIGH |
| 14 | 0.25 | 0.75 | 9.15 | HIGH/HIGH | HIGH | HIGH | HIGH |
| 15 | 0.26 | 0.74 | 8.08 | HIGH/HIGH | HIGH | HIGH | HIGH |
| 16 | 0.57 | 0 | 2.63 | LOW/LOW | LOW | LOW | LOW |
| 17 | 0.75 | 0 | 3.67 | LOW/LOW | LOW | LOW | LOW |
| 18 | 0.67 | 0.17 | 1.40 | LOW/LOW | LOW | LOW | LOW |
| 19 | 0.31 | 0 | 3.37 | LOW/LOW | LOW | LOW | LOW |
| 20 | 0.36 | 0 | 1.55 | LOW/LOW | LOW | LOW | LOW |
| 21 | 0.40 | 0 | 4.33 | LOW/LOW | LOW | LOW | LOW |
| 22 | 0.40 | 0 | 1.65 | LOW/LOW | LOW | LOW | LOW |
| 23 | 0.31 | 0 | 1.65 | LOW/LOW | LOW | LOW | LOW |
| 24 | 0.33 | 0 | 1.93 | LOW/LOW | LOW | LOW | LOW |
| 25 | 0.50 | 0.10 | 1.67 | LOW/LOW | LOW | LOW | LOW |
| 26 | 0.50 | 0 | 4.89 | LOW/LOW | LOW | LOW | LOW |
| 27 | 0.44 | 0.22 | 3.33 | LOW/LOW | LOW | LOW | LOW |
| 28 | 0.27 | 0 | 1.00 | LOW/NORMAL | LOW | LOW | LOW |
| 29 | 0.25 | 0 | 3.43 | LOW/LOW | LOW | LOW | LOW |
| 30 | 0.15 | 0.69 | 8.71 | HIGH/HIGH | HIGH | HIGH | HIGH |
| 31 | 0 | 1.00 | 8.20 | HIGH/HIGH | HIGH | HIGH | HIGH |
| 32 | 0 | 1.00 | 9.00 | HIGH/HIGH | HIGH | HIGH | HIGH |
| 33 | 0.09 | 0.91 | 8.82 | HIGH/HIGH | HIGH | HIGH | HIGH |
| 34 | 0.05 | 0.95 | 8.93 | HIGH/HIGH | HIGH | HIGH | HIGH |
| 35 | 0 | 1.00 | 10.00 | HIGH/HIGH | HIGH | HIGH | HIGH |
| 36 | 0.15 | 0.85 | 9.43 | HIGH/HIGH | HIGH | HIGH | HIGH |
| 37 | 0.17 | 0.83 | 10.00 | HIGH/HIGH | HIGH | HIGH | HIGH |
| 38 | 0 | 1.00 | 10.00 | HIGH/HIGH | HIGH | HIGH | HIGH |
| 39 | 0 | 1.00 | 10.00 | HIGH/HIGH | HIGH | HIGH | HIGH |
| 40 | 0.15 | 0.85 | 9.41 | HIGH/HIGH | HIGH | HIGH | HIGH |
| 41 | 0 | 1.00 | 10.00 | HIGH/HIGH | HIGH | HIGH | HIGH |
| 42 | 0 | 1.00 | 10.00 | HIGH/HIGH | HIGH | HIGH | HIGH |
| 43 | 0 | 1.00 | 10.00 | HIGH/HIGH | HIGH | HIGH | HIGH |
| 44 | 0.29 | 0 | 1.35 | LOW/LOW | LOW | LOW | LOW |
| 45 | 0.44 | 0 | 1.63 | LOW/LOW | LOW | LOW | LOW |
| 46 | 0.56 | 0 | 1.92 | LOW/LOW | LOW | LOW | LOW |
| 47 | 0.67 | 0 | 5.63 | LOW/LOW | LOW | LOW | LOW |
| 48 | 1.00 | 0 | 4.50 | LOW/LOW | LOW | LOW | LOW |
| 49 | 0.53 | 0 | 3.20 | LOW/LOW | LOW | LOW | LOW |
| 50 | 0.69 | 0.06 | 1.91 | LOW/LOW | LOW | LOW | LOW |
| 51 | 0.38 | 0 | 2.44 | LOW/LOW | LOW | LOW | LOW |
| 52 | 0.40 | 0 | 2.81 | LOW/LOW | LOW | LOW | LOW |
| 53 | 0.67 | 0.17 | 4.93 | LOW/LOW | LOW | LOW | LOW |
| 54 | 1.00 | 0 | 1.17 | LOW/LOW | LOW | LOW | LOW |
| 55 | 0.40 | 0 | 7.00 | LOW/HIGH | LOW | LOW | LOW |
| 56 | 0.29 | 0 | 1.39 | LOW/LOW | LOW | LOW | LOW |
| 57 | 0.43 | 0 | 2.90 | LOW/LOW | LOW | LOW | LOW |

1 The nucleus detector detects and classifies nuclei image-wide in a FISH image
2 The signal detector detects, classifies and counts FISH signals in each nucleus detected by the first RetinaNet pipeline
3 Ratio-1 represents number of low-grade nuclei divided by the number of all classified nuclei. A value greater than 0.2 indicates LOW stage of the FISH image.
4 Ratio-2 represents number of high-grade nuclei divided by the number of all classified nuclei. A value greater than 0.4 indicates HIGH stage of the FISH image.
5 The average of all detected HER2/CEN17 ratios among all nuclei in one FISH image. A value greater than 1.0 indicate LOW stage and greater than 6.0 HIGH stage of the FISH image.

## Conclusions

In this study, we developed a deep learning pipeline for analyzing Fluorescence *in situ* hybridization (FISH) images regarding the image-wide detection of interphase nuclei and their classification depending on the HER2 gene amplification level. The pipeline can be useful in assisting pathologists in analyzing the HER2 gene amplification stage of a breast cancer or gastric cancer samples by automatically analyzing high quality FISH images. Image-wide ratios representing the number of abnormal nuclei in relationship to all classified nuclei are calculated which serve as guideline for classifying the HER2 gene amplification status of the corresponding tumor sample from which the FISH image originated from. It can also be used for automatic investigation in retrospective studies of large amounts of documented FISH images collected over several years for re-evaluation. Another application could be the enhancement of the documentation quality of the images. Furthermore, an anonymized and human-independent evaluation of the HER2 gene amplification level is possible. Analyzing one FISH image including the generation of the annotated image data and the report occurs in less than a second which is orders of magnitude faster compared to human visual evaluation and manual report generation. Therefore, we interpret our pipeline as a first step towards the automation of the HER2 gene amplification detection in FISH images.

Our pipeline consists of the successive application of two state-of-the-art CNNs for localization and classification (nucleus detector and signal detector). In the first step the nucleus detector detects and classifies nuclei in the FISH images and calculates two ratios: ratio-1 represents how frequently nuclei with a low amplification status of HER2 genes occurred and ratio-2 indicates the same for nuclei with high amplification status of the HER2 gene. In the second step, the signal detector, identifies and classifies FISH signals for each nucleus into HER2 signals, CEN17 signals and HER2 cluster, which consists of multiple, non-distinguishable HER2 signals. An average HER2/CEN17 image-wide ratio is calculated on the basis of all nucleus-specific HER2/CEN17 ratios and serves for decision making on the image-wide HER2 gene amplification status of the FISH image. The reliability of our pipeline was demonstrated to be comparable to the pathologist with an accuracy of 96% based on 57 annotated FISH images (55 out of 57 images). Apart from its accuracy, our two-step pipeline provides additional robustness by classifying every nucleus twice. Moreover, the detailed reports at the image level as well as at the level of an individual nucleus provide important information that largely increases the interpretability of the automated detection of the HER2 amplification status. One limitation of this approach is need for annotation of bounding boxes and classification of nuclei and FISH signals. This requires considerable addition work as compare to an alternative approach where image-wide classification of HER2 amplification status is directly learned from FISH whole slides. Indeed, it is possible to train off-the-shelf networks such as ResNet^23^ or Inception^33^ to achieve comparable accuracy as our pipeline. However, this strategy completely lacks the AI interpretability which we consider an essential requirement for clinical applicability. Our deep learning pipeline will only be applied in clinical practice and involved in decision about treatment options if pathologists are able to understand how the system arrives at a particular decision.

Software-based systems for automated detection of HER2 gene amplification status in breast cancer samples have already been developed^34–36^, all based on conventional image analysis algorithms. These software solutions are strictly dependent on accurate preparation of the FISH slides, which otherwise could not be well analyzed by the software pipeline. In contrast, our pipeline can easily be trained under laboratory specific conditions and is not dependent on strict preparation conditions and therefore is highly flexible to be used in any laboratory under laboratory conditions. The conventional image analysis pipelines achieve a high level of agreement with the pathologist (e.g. acc=96%-99% and κ=0.89 or κ=0.94^36^ or acc=96%-97% and κ=0.92 to κ=0.94^34^ (depending on categorization strategy)) on the single sample level which is comparable to our success rate of 96% accuracy on a FISH image-wide level. However, since no quantitative data is available on their performance at the level of a single nucleus, our results cannot be directly compared to these previous studies. At this level, our results show that our system achieves agreements that correspond to those of a team of pathologists.

However, there are still limitations of our pipeline: An important issue is the difficulty in predicting the HER2 gene amplification status in FISH images of relatively low quality characterized by high background noise, low signal-to-noise ratio, a large number of artifacts, strong differences in nuclei shape, weak signals, or truncated/overlapping nuclei. To overcome these limitations one would need to either (1) greatly enlarge the manually annotated training data set by incorporating many thousands of corresponding low-quality examples, (2) improve the image acquisition practice in clinical routine to obtain higher quality images, or (3) further increase the number of pathologists that provide annotations. However, due to performing the FISH diagnostics on slides originating from FFPE material it might be difficult to obtain higher quality images. FISH slides originating from FFPE samples are of much lower quality compared to slides from fresh material. Training our pipeline on these samples would largely enhance its performance on future cases of similar reduced image quality. Nevertheless, even in its current form our pipeline (trained on high quality FISH images) makes surprisingly accurate predictions on the HER2 amplification status of the tumor on these low quality FISH images demonstrating the general potential of deep learning on this task (**Suppl. Fig. 1**). It should be noticed that, in clinical practice, pathologists do not analyze every nucleus in a FISH image. Instead, a certain number of nuclei (at least 20) are pre-selected and all other nuclei that are difficult to analyze (e.g. due to low image quality) are excluded. Additionally, variations in the experimental setup among different pathology labs might result in different shape, structure and nuclei composition of the FISH images (e.g. used antibodies and fluorophores, tissue type, tissue preparation protocol, consideration of DAPI staining, fluorescence microscope type and parameters). Therefore, a customization of our pipeline, e.g. setting different thresholds, and additional training of both networks will be necessary to adapt the detection and classification pipeline to lab-specific conditions and lab-specific investigated tissue types in order to automatize the HER2 amplification detection of tumors in other pathology labs.

Pathologists normally analyze the FISH slides directly under the fluorescence microscope by acquiring only a single 2D image at a specific focus position of the microscope. Consequently any volumetric image context that otherwise could provide additional classification information is lost. Therefore, a deep learning pipeline based on nuclei detection and classification on a stack of images representing the 3D volume of the FISH slide is likely to show improved accuracy compared to the 2D solution used in our study. Additionally, future solutions could directly implement one-stage detectors or similar CNN architectures into the fluorescence microscope for instantly classifying the nuclei while the pathologist is observing it. A comparable solution was recently developed by Google Inc. for marking tumor areas in Hematoxylin and Eosin stained slides^37^. Alternatively, a fully automated software solution recording all layers and positions of a FISH slide as volumetric input to the deep learning-based nuclei detection and classification pipeline might be used.

## Authors’ contributions

FZ, WdB and PH wrote the manuscript. FZ, WdB and PH designed the study. FZ, WdB, MW, RM and TW planed and conducted the bioinformatics data preparation and analysis. SZ and DEA generated the FISH data. KF, SZ, CS, PH, DEA and GB performed the pathological analysis of the data. KF, DEA, IR and GB supervised the project and assisted in the writing of the manuscript.

## Acknowledgments

We thank the Machine Learning Community (MLC) Dresden and Uwe Schmidt (CSBD) for helpful input, comments and inspiration. We also thank Regina Pohlers and Ines Kaiser for technical assistance in generating FISH images and annotating the ground-truth FISH data.

---

**Supplemental Figure 1.**
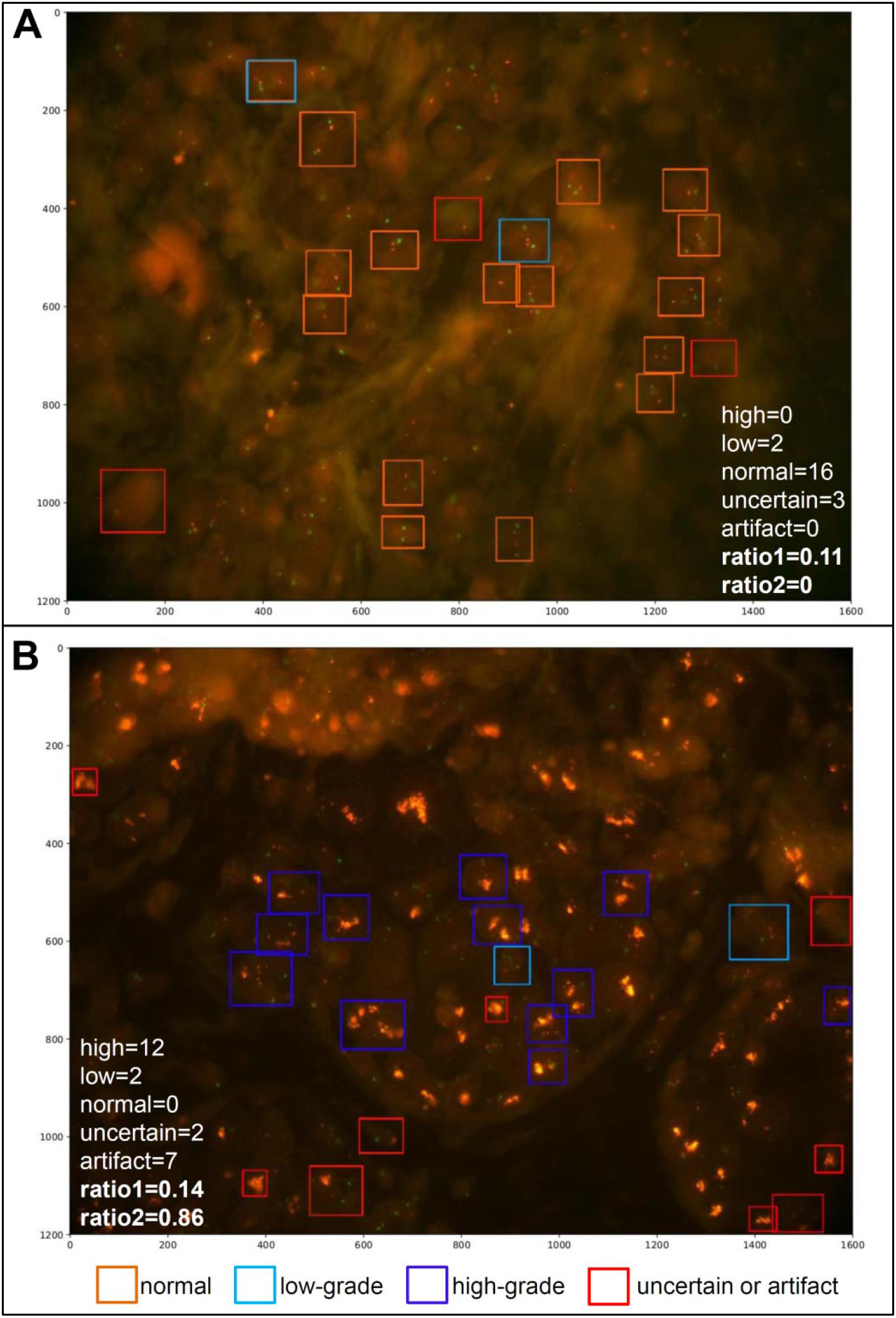
Two examples of the application of our pipeline on FISH images of low quality. In **(A)** a normal stage was detected and corresponds to the decision of a pathologist. In **(B)** a high-grade stage was detected which also corresponds to the pathologists decision on the FISH image. Numerous nuclei have not been detected in both images indicating the limitations of our system on FISH images of very low quality. Training on a large set of FISH images of low quality would enhance the accuracy in detecting most nuclei.

