## Supplementary figures and images for "Automated detection of the HER2 gene amplification status in Fluorescence *in situ* hybridization images for the diagnostics of cancer tissues"

### Supplemental Figure 1

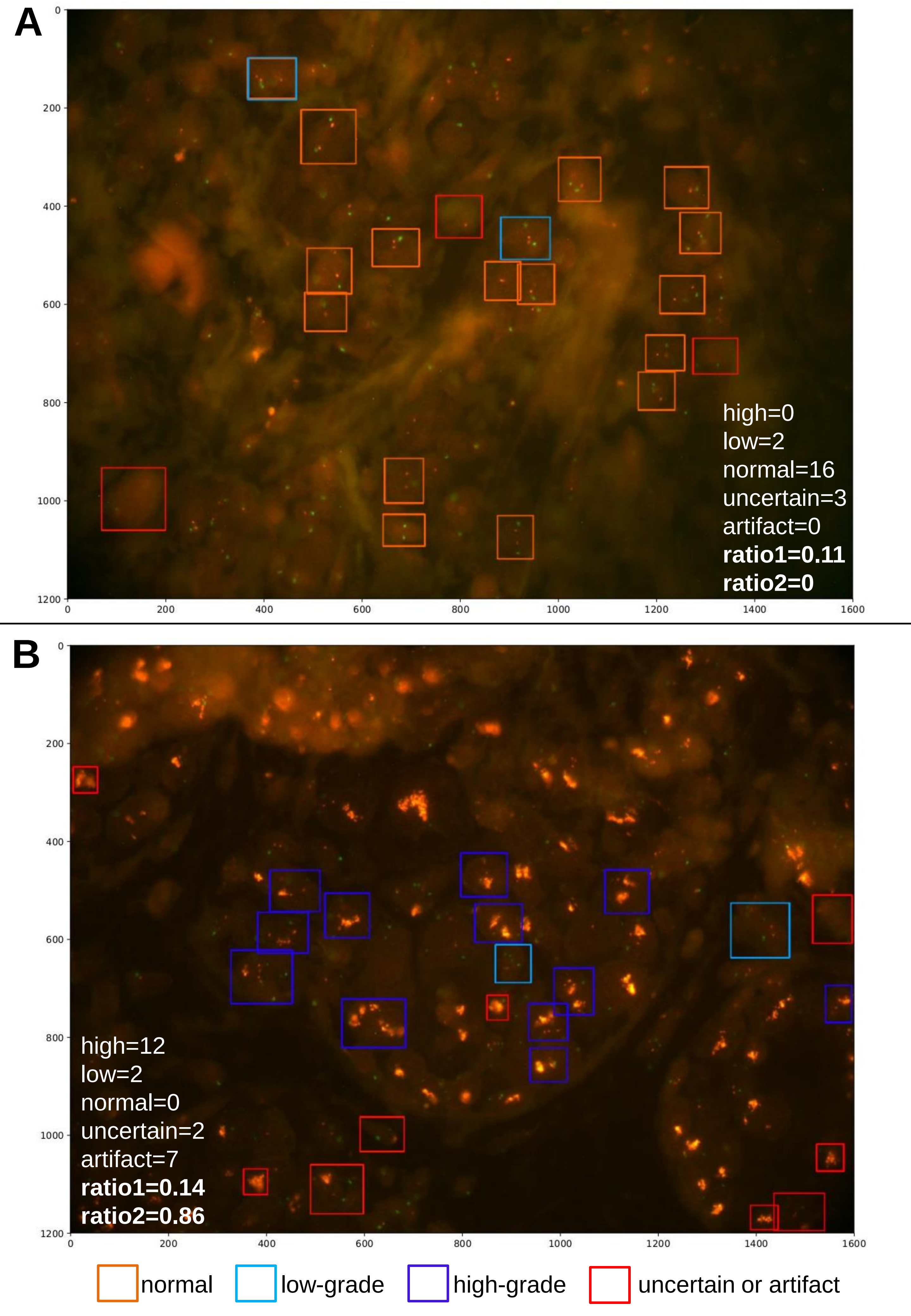
